# Neurodegeneration exposes firing rate dependent effects on oscillation dynamics in computational neural networks

**DOI:** 10.1101/663187

**Authors:** D. Gabrieli, Samantha N. Schumm, B. Parvesse, D.F. Meaney

## Abstract

Traumatic brain injury (TBI) can lead to neurodegeneration in the injured circuitry, either through primary structural damage to the neuron or secondary effects that disrupt key cellular processes. Moreover, traumatic injuries can preferentially impact subpopulations of neurons, but the functional network effects of these targeted degeneration profiles remain unclear. Although isolating the consequences of complex injury dynamics and long-term recovery of the circuit can be difficult to control experimentally, computational networks can be a powerful tool to analyze the consequences of injury. Here, we use the Izhikevich spiking neuron model to create networks representative of cortical tissue. After an initial settling period with spike-timing-dependent plasticity (STDP), networks developed rhythmic oscillations similar to those seen *in vivo*. As neurons were sequentially removed from the network, population activity rate and oscillation dynamics were significantly reduced. In a successive period of network restructuring with STDP, network activity levels were returned to baseline for some injury levels and oscillation dynamics significantly improved. We next explored the role that specific neurons have in the creation and termination of oscillation dynamics. We determined that oscillations initiate from activation of low firing rate neurons with limited structural inputs. To terminate oscillations, high activity excitatory neurons with strong input connectivity activate downstream inhibitory circuitry. Finally, we confirm the excitatory neuron population role through targeted neurodegeneration. These results suggest targeted neurodegeneration can play a key role in the oscillation dynamics after injury.

**Author Summary:** In this study, we study the impact of neuronal degeneration – a process that commonly occurs after traumatic injury and neurodegenerative disease – on the neuronal dynamics in a cortical network. We create computational models of neural networks and include spike timing plasticity to alter the synaptic strength among connections as networks remodel after simulated injury. We find that spike-timing dependent plasticity helps recover the neural dynamics of an injured microcircuit, but it frequently cannot recover the original oscillation dynamics in an uninjured network. In addition, we find that selectively injuring excitatory neurons with the highest firing rate reduced the neuronal oscillations in a circuit much more than either random deletion or the removing neurons with the lowest firing rate. In all, these data suggest (a) plasticity reduces the consequences of neurodegeneration and (b) losing the most active neurons in the network has the most adverse effect on neural oscillations.

## Introduction

Traumatic Brain Injury (TBI) is a prominent cause of disability in the US (1). Perhaps because of growing awareness of the consequences of TBI, emergency department visits for TBI have increased 47% from 2007 to 2013 (2). Although many of these injuries produce no long-term deficits, a fraction of injuries produce cognitive and psychological impairments that can last years after the original insult (3–5). As a result, more than 5 million people in the US live with significant consequences of TBI, contributing to an estimated 70 billion dollars annually in medical and non-medical costs (6). Effective recovery from TBI remains a challenge because no two injuries are exactly alike, leading to a unique injury and recovery pattern for each TBI patient.

One key feature in TBI recovery is how the structural and functional networks in the brain evolve over time after injury to guide the cognitive recovery processes (7–11). Recent studies have shown alterations in brain circuitry after moderate and severe injury affect the coordination among functional brain networks (12). With the development of models to predict the overall changes in brain networks during different tasks, there is an emerging consensus that the dynamic network that connects different brain regions can influence cognitive and psychological alterations after injury (13–16). However, the unique circumstances that cause each TBI make it difficult to predict which injuries will likely lead to long-term changes in brain function. These challenges exist especially at the cellular scale, where the neuronal degeneration that can occur days to weeks after a TBI can affect the function of local microcircuits throughout the brain (17,18).

To this end, computational models can be a useful tool to understanding how neuronal damage can ultimately contribute to the impairments in circuit function after TBI. In general, these models can account for the mechanisms of acute injury to the network (e.g., primary axotomy, membrane permeability changes, receptor dysfunction) and secondary changes that can also trigger neuronal loss (18–22). Despite the many experimental methods exist to explore neural activity at different scales – e.g., single unit recording, local field potential recordings representing the aggregate activity of neuronal ensembles, and high speed calcium imaging to explore neuronal activation in awake animals – these techniques are difficult to develop this precise relationship between neurodegeneration and network dynamics (23–26). Alternatively, computational models can systematically examine the effect of damaging neurons within an integrated network without the influence from variable upstream circuitry. We can gain critical information that would be impossible or impractical to acquire using conventional methods.

Building upon past studies that examined how neuronal connectivity and injury patterns can lead to activity patterns which resemble posttraumatic epilepsy (27,28), we use a computational model to examine the effect of neurodegeneration on the spontaneous activity of neural circuits. We utilize microcircuits that resemble isolated cortical circuitry to identify the exact relationship between local changes in network function and degeneration without the complexity of large-scale interconnected topology. We focused our work on how spike-timing dependent plasticity affected the recovery of these circuits after degeneration and how degeneration in populations of neurons can play specific roles in altered network function. Together our results demonstrate how neurodegeneration affects the dynamics of a microcircuit, and the importance of spike timing dependent plasticity in repairing damaged microcircuits after injury.

## Results

To date, there is no clear evidence in the literature that suggests neurons with specific topological properties are more vulnerable than others to traumatic injury, although there are indications that specific regions of anatomic structures (e.g., CA3 in the hippocampus, cingulate, or thalamus) that may preferentially show neuronal damage (29–31). We first examined how the random deletion of neurons affected the pattern of neuronal activity and oscillations in our network. Without any neuronal deletions, our networks showed an average firing rate of 4.7 ± 0.1 Hz and an average oscillation frequency of 12.4 +/ − 0.4 Hz (Figure 1A-C). This activity level and the presence of oscillations are consistent with past computational models using similar methods (32–34).

**Figure 1.**
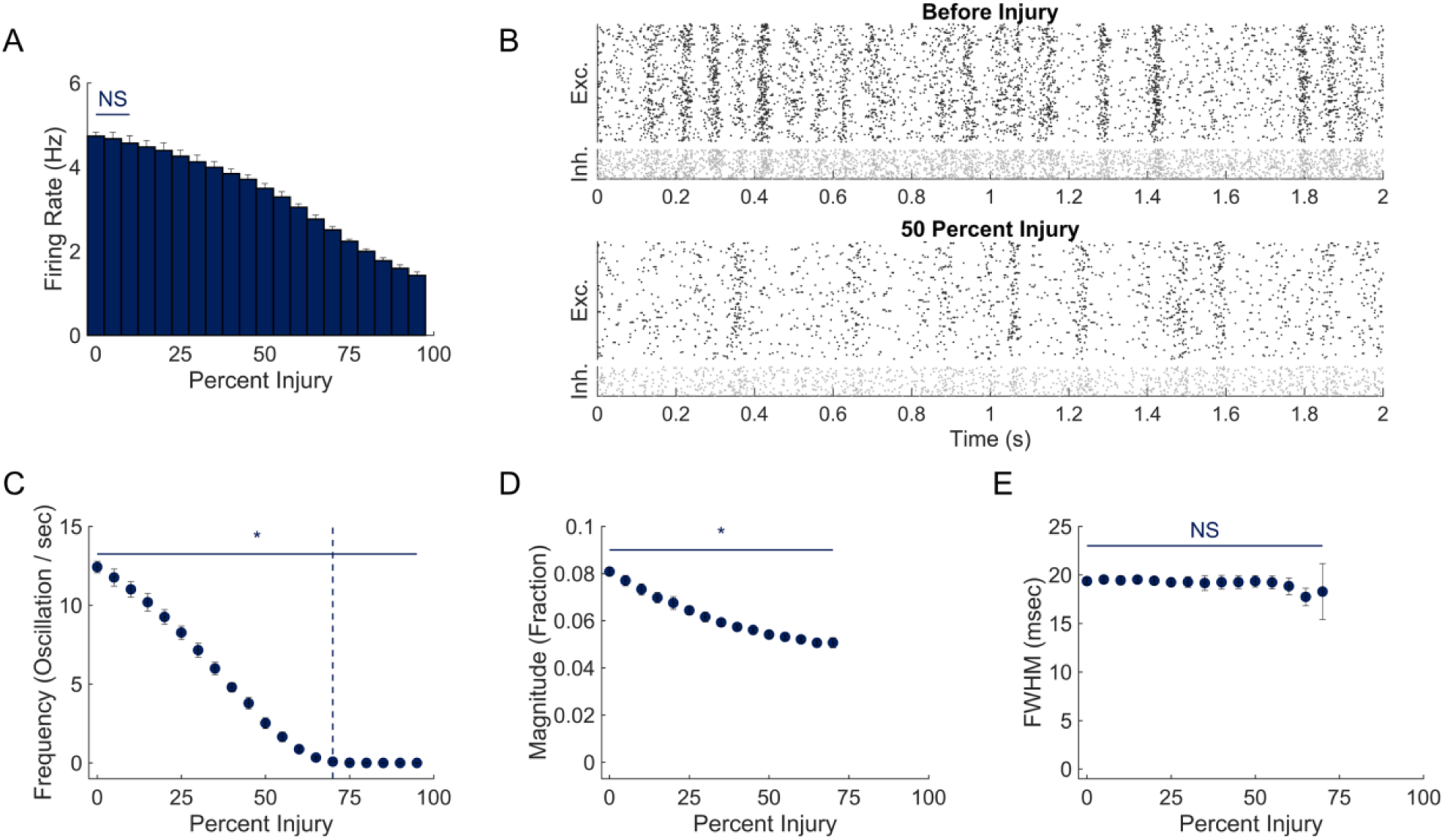
Effect of Neurodegeneration on Network Dynamics. **(A)** Network firing rate was significantly reduced after 15% injury and continued to decline with further damage. **(B)** Representative raster plots before and after 50% injury. **(C**,**D)** Network oscillation frequency and magnitude changed significantly from baseline at 5% injury. Network oscillations were not observed after 85% injury. Oscillations were not present in simulations with greater than 70% injury, demarked by the vertical dashed line. **(E)** Random neurodegeneration did not significantly impact oscillation FWHM.

Patterns of activity and oscillations slowly changed with the progressive deletion of neurons in the network. The average firing rate significantly decreased when deleting 15% or more of the network neurons (One way ANOVA with Tukey Kramer post-hoc P<.001; Figure 1A). At the peaks in oscillation activity, we found that 8.1 ± 0.2 % of the neuronal network was activated at baseline (Figure 1D). Random removal of neurons also led to a significant decrease in oscillation frequency and oscillation magnitude, with these changes appearing for lower damage levels (>5% or more deleted neurons (One way ANOVA with Tukey Kramer post-hoc; p=0.001 and p<0.001, respectively; Figure 1C, D). The duration of an oscillation was most resistant to neuronal loss, requiring at least 80% neuronal loss to show a significant decrease (One way ANOVA with Tukey Kramer post-hoc; Figure 1E).

We next considered if STDP would repair the functional deficits appearing in networks after damage. The average firing rate of neurons in the network increased significantly over a broad range of damage when STDP rebalanced synaptic weights (20-80%; Figure 2A; representative changes in Figure 2B). At lower levels of damage (5-60%), average neuronal activity was not significantly different from undamaged networks (Figure 2A). Similarly, plasticity allowed a significant increase in oscillation frequency and magnitude relative to networks in which the synaptic weight was held constant (Figure 2C, D, E). Unlike average firing rate, though, plasticity did not recover the oscillation frequency, magnitude or duration to levels observed in undamaged networks. Beyond 25% damage to the network, oscillation width significantly increased relative to undamaged networks (One way ANOVA with Tukey Kramer post-hoc, Figure 2E).

**Figure 2.**
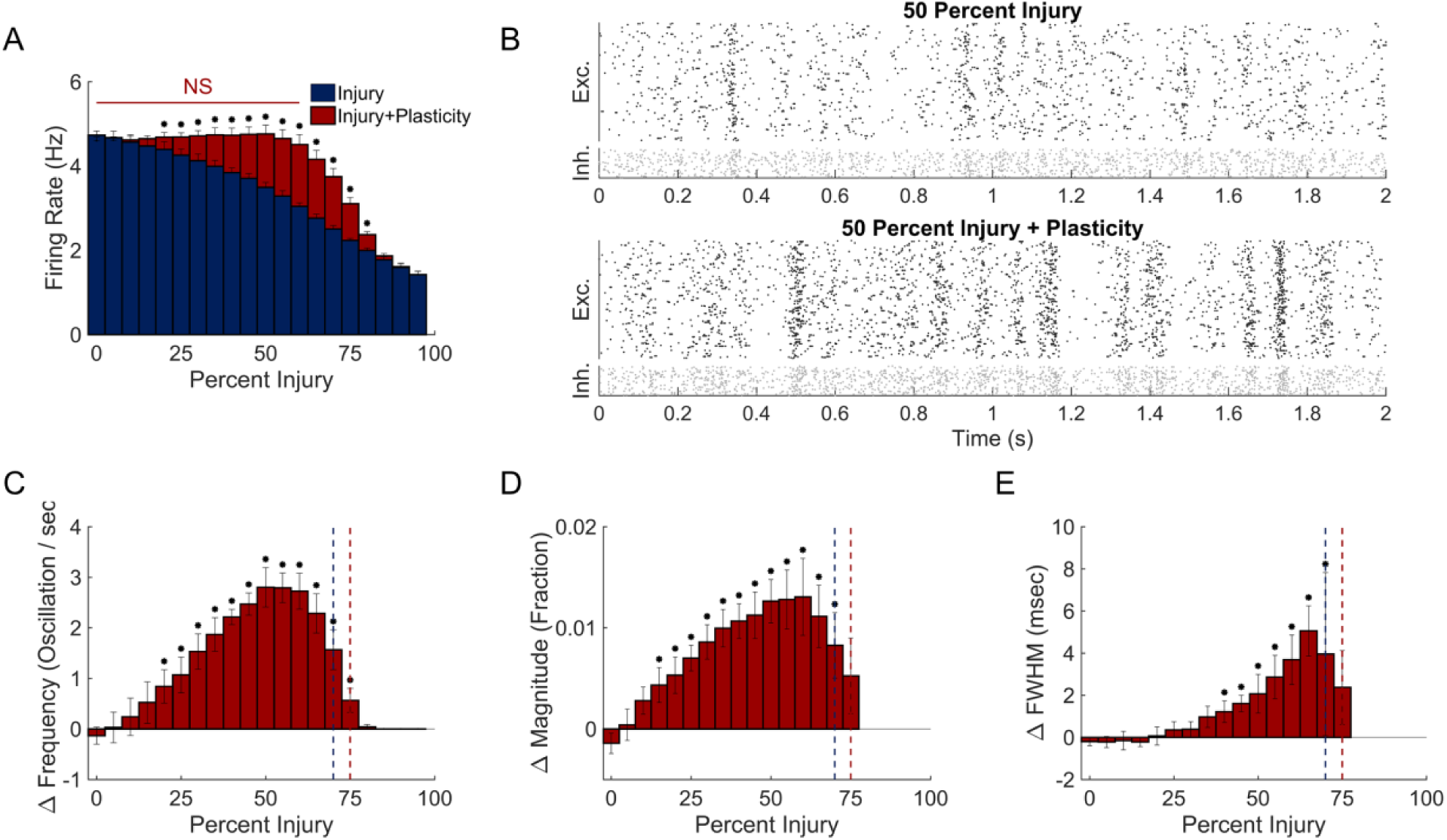
Role of Plasticity in Network Recovery. **(A)** Four hours of recovery with STDP fully restored neuron activity back to baseline up to 60% damage. **(B)** Raster plots after 50% neurodegeneration and subsequent STDP recovery. **(C**,**D)** Network oscillation frequency and magnitude recovered significantly at moderate levels of damage (two sample t-test with Bonferroni correction for multiple comparisons). These changes were not sufficient to return to baseline, but partially restored function. Dotted vertical line at 70% and 75% injury denote the last injury that all simulations contained oscillations in pre- and post-plasticity simulations respectively. **(E)** FWHM differences after plasticity were significantly different from baseline and significantly different from immediately post-injury in the range of 40-70% injury.

After observing that spike-timing dependent plasticity helped introduce resilience to damage, we next explored if there were specific connectivity features of individual neurons that influenced or explained part of this resilience. For each neuron in an undamaged network, we computed a neuron connectivity index as the normalized difference of total synaptic input strength and the total output strength. This neuron-connectivity index correlated with the average neuronal firing rate; neurons with high index showed higher firing rates than neurons with low index (Figure 3A). The relationship between input/output strength and firing rate was stable while allowing the synaptic weights to adjust via STDP over 4 simulation hours. In addition, these relationships did not change across a broad range of neuronal network parameters that included synaptic strength, neuron parameters (a-d) that could change neuronal type (33), and connection number among neurons in the network.

**Figure 3.**
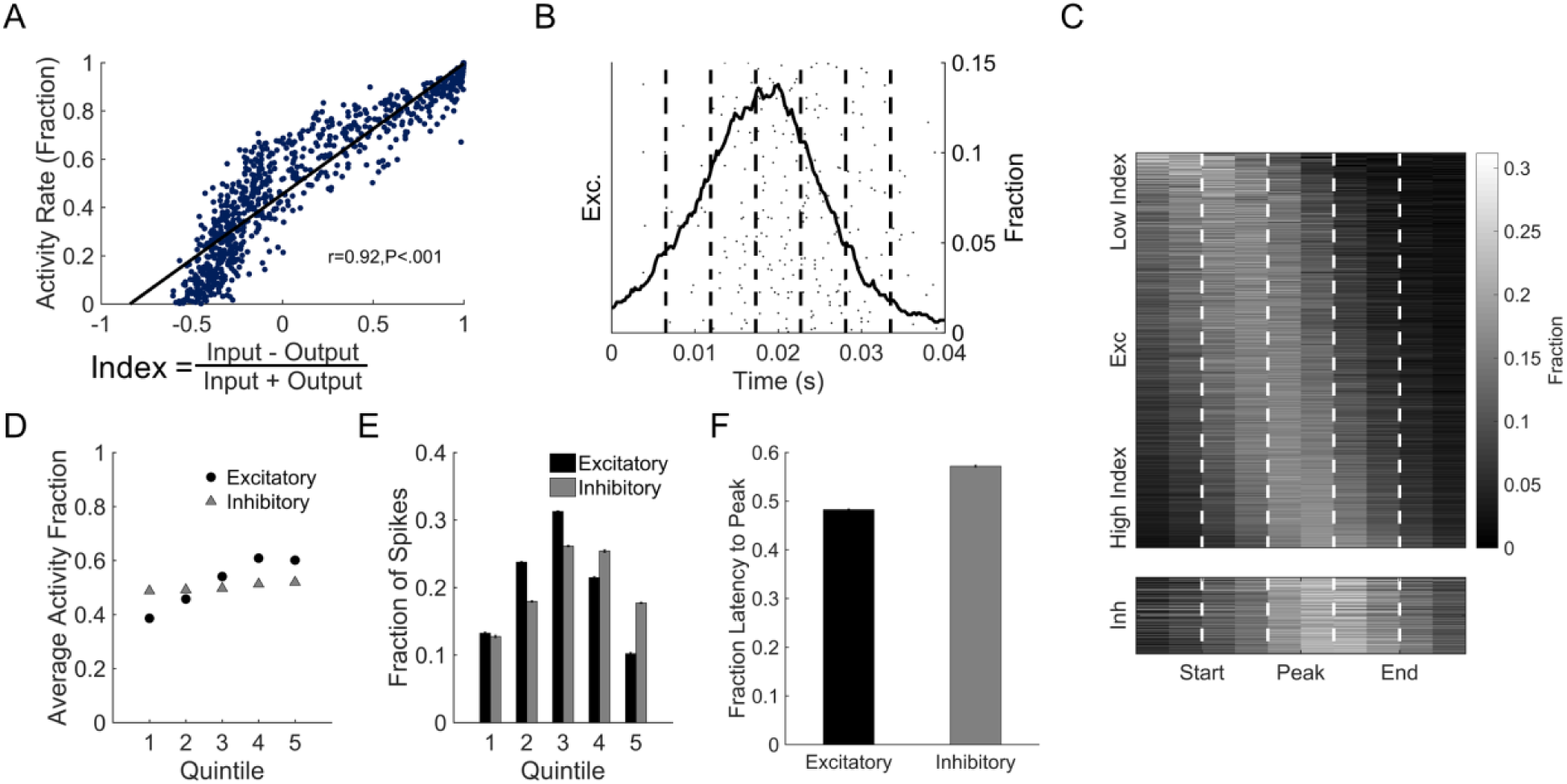
Activity Dependent Functional Roles of Neurons. **(A)** After initial settling, networks developed structure-function relationships. We defined the network connectivity index as the difference between input and output strengths normalized to total strength. Firing rate was significantly correlated with network connectivity index. **(B)** To test the roles of neurons with different activity profiles in oscillations, we divided each oscillation into ten equal segments centered around the peak activation period. We then tracked when each neuron fired within the oscillation. **(C)** Low index neurons tend to fire early in network oscillations, suggesting a role in oscillation initiation. High index neurons conversely fire at the peak of the oscillation and activate downstream inhibitory circuitry to stop oscillatory behavior. **(D)** Low activity excitatory neurons were more likely to be active early in oscillations. Later stages showed increased activation of highly active excitatory neurons. Inhibitory neurons showed no activity dependence in their firing time within oscillations. **(E-F)** Excitatory neurons had increased activation to peak oscillation magnitude, with decreasing activation after the peak. Inhibitory neurons had delayed activation, primarily firing at peak or late in the oscillation.

We then sought to find if either the initiation or termination of the oscillations were related to neuronal activity and, in turn, connectivity strength. Using our definition of the beginning and end of an oscillation (see Methods; representative oscillation appears in Figure 3B, C), we divided an oscillation period into quintiles. In general, excitatory neurons with low inputs relative to their outputs (i.e., low index) fired primarily during the initiation period of an oscillation, while high input strength excitatory neurons fired near the peak of an oscillation and activated inhibitory neurons to arrest the oscillation. The average firing rate of excitatory neurons within each of the first four quintiles significantly increased, while the firing rate significantly decreased in the fifth quintile (One way ANOVA with Tukey Kramer post-hoc (p<.001; Figure 3D)). In comparison, we could not identify any dependence on firing time and oscillation period for inhibitory neurons (Figure 3D). Excitatory neurons most commonly fired during the peak of the oscillation, decreasing on both sides of peak (Figure 3E). Inhibitory neurons had a preference to fire later in the oscillation, peaking between the 3^rd^ and 4^th^ quintile (Figure 3E). Together, these results indicate that the initiation of an oscillation corresponds with the activations of neurons with low firing rates, while the termination of the oscillation begins with the simultaneous recruitment of neurons with high firing rates and activating a significant fraction of the inhibitory neurons.

As excitatory neurons with specific connectivity and activity patterns correlated with oscillation dynamics, we next considered if the targeted deletion of either activity type would alter the neural circuit dynamics. Using the random deletion of neurons as a comparison, we explored the change in functional network characteristics that would occur if we progressively deleted the neurons with the lowest average activity. With this strategy, we found that average activity rate would significantly decrease when damage exceeded 5% of the network (Figure 4A-B). Similar to random deletion, less damage was required to significantly changes oscillation frequency (Figure 4C; damage > 5%) than activity rate. The deletion of the lowest firing rate neurons additionally led to significant change in the average width of an oscillation (Figure 4E). Plasticity returned the average activity in the network to baseline up to 15% damage; beyond this level, plasticity did improve average firing rate up to 90% injury (Figure 4A). In comparison, plasticity improved the oscillation frequency (and magnitude) over a range of the injury levels (10-65% and 5-55% respectively) but did not reach baseline levels.

**Figure 4.**
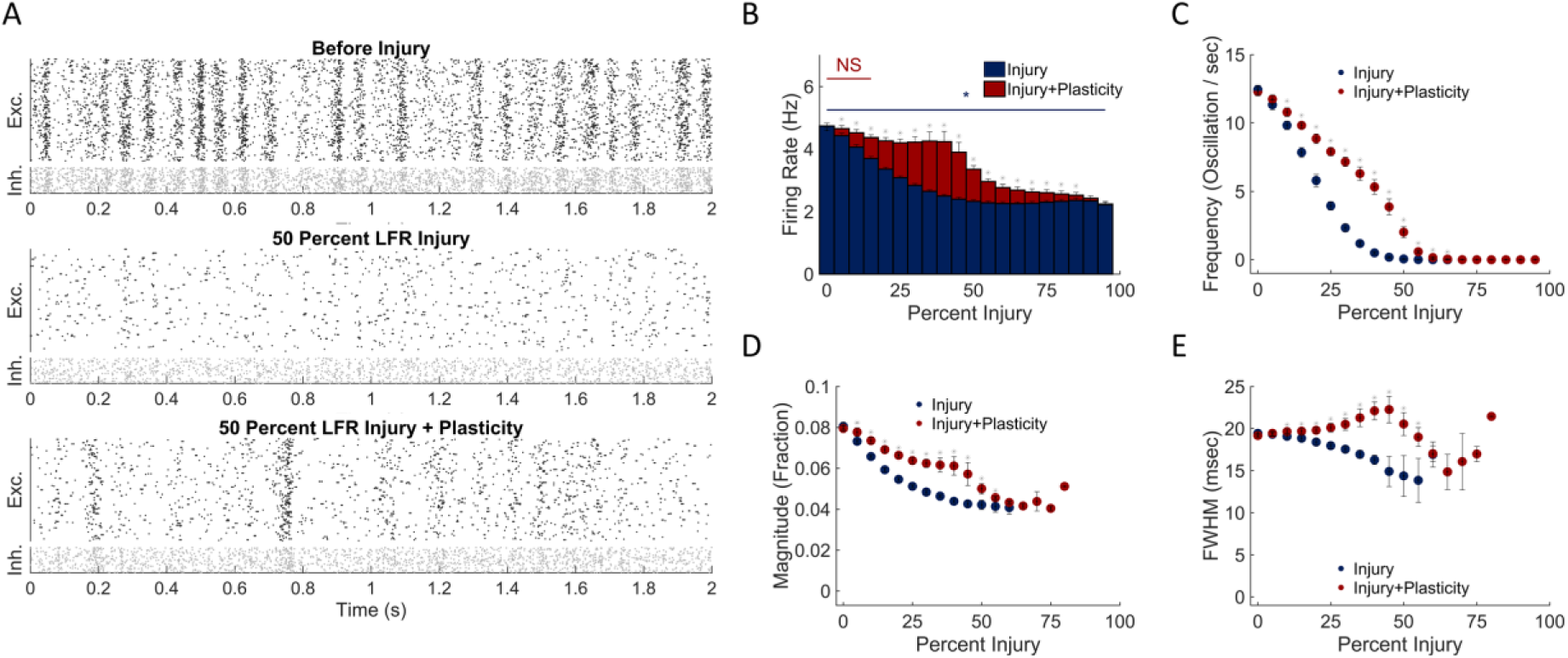
Effect of deleting relatively inactive neurons is partially recovered with plasticity. **(A)** Representative raster plots before injury, at 50% low firing rate (LFR) excitatory neuron injury, and after network restructuring with STDP. **(B)** LFR damage produced a rapid decline to baseline firing after injury that significantly recovered with STDP. **(C**,**D)** This restoration was primarily due to a recovery of network oscillation frequency and magnitude. **(E)** Changes in oscillation width occurred relative to baseline, primarily following damage exceeding the threshold of full firing rate restoration.

In contrast to these results, the progressive deletion of neurons with the highest activity rate did not significantly change the overall average activity of the network until more than 20% of neurons were removed (Figure 5A-B). Removing the neurons with highest activity led to a progressive decrease in oscillation frequency that was significantly different than deleting the same fraction of low firing rate neurons (t-test with Bonferonni correction for multiple comparisons; p<.001; Figure 4C & 5C). Unlike the random deletion of neurons, the width of oscillations increased significantly over a broad injury range (Figure 5E; 15-55%). While the oscillation frequency and/or magnitude changed with this injury approach, plasticity only modestly affected the oscillation frequency, magnitude and average width at higher damage levels. (Figures 5C-E).

**Figure 5.**
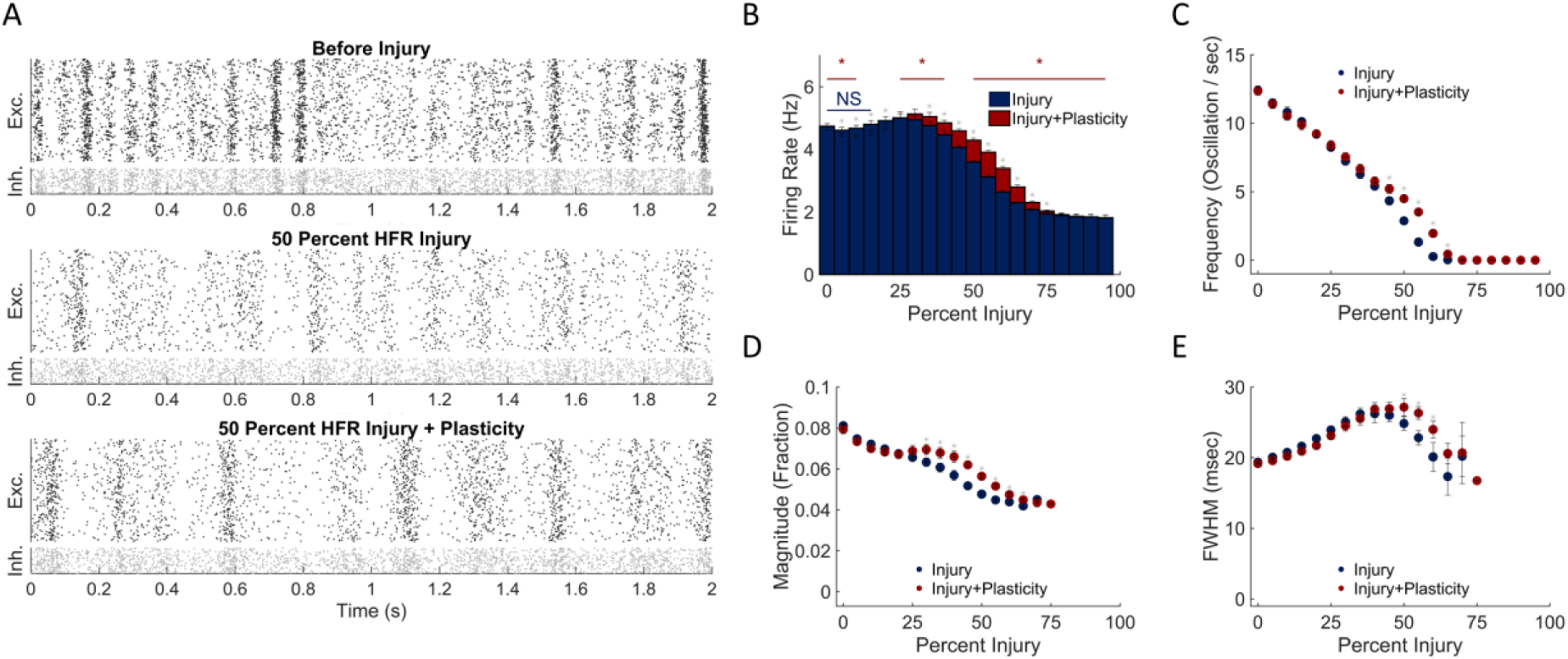
Removing highly active neurons predominantly affects oscillation dynamics. **(A)** Raster plots before injury, at 50% high firing rate (HFR) excitatory neuron injury, and after network restructuring with STDP. **(B)** Network activity did not significantly vary from baseline after HFR neurodegeneration. **(C-D)** Oscillation frequency and magnitude after plasticity were significantly different at the same injury level at higher levels of injury. **(E)** Oscillation width increased above baseline levels after damage and was maintained following restructuring with STDP.

## Discussion

Here, we modeled the effects of neurodegeneration on the functional dynamics of neural circuitry. We utilized an established spiking neural network model to investigate how damage and plasticity impact network firing and oscillatory behavior. We show that neurodegeneration, regardless of whether it was random or based on neuronal firing rate, significantly decreased network activity and oscillation dynamics. For all types of neurodegeneration, activity was significantly restored with spike-timing dependent plasticity. Deleting highly active neurons led to a marked increase in oscillation duration, while deleting neurons with low activity reduced the initiation of oscillations without decreasing their frequency. These results suggest that the degeneration or inactivation of specific neuron activity profiles can differentially affect oscillation dynamics of neuronal circuits.

We used several simplifying assumptions to examine the potential impact of neurodegeneration. First, we use generalized topologies and neuron spiking behaviors of cortical neurons based on simplistic rules developed from observations *in vivo* (28,32,33,35,36). We recognize that neurons within specific brain regions can vary greatly in the connectivity preferences and spiking behavior (37–39), and therefore some of our observations on neuronal degeneration may not apply universally to all brain regions. We did explore if our observations were influenced by different distance-dependent connection algorithms and found that the general decline in activity and the restorative effect of plasticity did not depend on the initial spatial connections in the network. Although we anticipate that our general findings could inform predictions for networks of more complex, region specific topologies, many of the regions commonly damaged in TBI (e.g., hippocampus, thalamus, and cingulate (29,30)) do not have clear estimates of neuronal connectivity. Once available, we envision creating specific computational network models to assess the dynamics would differ with deletion in these specific regions.

Second, we used Izhikevich integrate-and-fire neuron model to approximate neuron activity, which may not fully represent the complexity of real circuitry. Because the Izhikevich model accurately predicts neuronal spike times for a variety of neuron spiking behaviors (33,40,41), this model should be representative of *in vivo* neural activity for the metrics we utilize in this study. Extending the current work into a more computationally complex neuron model with higher-order biological features (reviewed in (42)) could provide additional information into the timing of activation and synaptic inputs across an individual neurons (43). However, these changes would likely affect all neurons in the network similarly and will not significantly impact the overall estimates of activity and oscillatory behavior.

Finally, we considered whether the broad distribution of neuronal types employed in larger scale Izikevich integrate-and-fire models were important to include in this study. However, without any insight into how more homogeneous circuits remodel in response to trauma, we believed this additional complexity would prevent one from gaining clear insight into which types of neuronal deletion would be particularly damaging to circuit dynamics.

In general, our observed neural activity patterns match the general spiking patterns and network wide oscillations in computational networks of similar size (e.g., (27,33)), and even more complex networks of much larger size and complexity (32). One key characteristic we observed in our simulations was the coordinated activation of a subset of neurons in regular periodic intervals over the entire simulation period. Again, these waves of neuronal activation are distinct from a near simultaneous activation or bursting of the network that can appear in some studies (27). These oscillations of neuronal activity, where 10-15% of the network was activated, were the first to show a significant decrease in oscillation frequency after neurons were deleted from the network. Although neuron characterization traditionally depends on functional spiking properties of neurons (44–46), our results show that plasticity is an important mechanism to produce functional diversity. Moreover, our results identified subpopulations of neurons that could either help trigger or suppress these periods of high network activity. To our knowledge, showing that neurons with low activity rates preferentially activate oscillations in a network has not been reported in past modeling studies, nor are we aware of past reports showing that neurons with high firing rates are important for quieting periods of high activity. Together, these data suggested that targeting of specific neuronal populations for degeneration would preferentially affect neuronal dynamics, a prediction we confirmed with subsequent simulations.

The rhythmic oscillations may also have important consequences on the synchronization, or coherence, of activity across brain regions. Coherence among neuron populations has been shown to be important for attention and memory (47–49), cognitive processes that are commonly affected after traumatic brain injury. Selective degeneration of low firing rate neurons reduces the likelihood of initiating an oscillation, in turn lowering the coherence with other brain regions downstream of the injured microcircuit. Therefore, losing this neuronal subpopulation would appear to play a significant role affecting information relayed across brain regions, an aspect of information processing that has appeared in network-based studies of TBI (13,50). In comparison, losing neurons with high firing rates would appear to have less consequence on the coordinated oscillations among regions because these neurons would not affect the emergence of an oscillation, and only slightly lengthen the duration of an oscillation. However, we cannot completely discount the impact of losing high firing rate neurons because the lengthening of a specific oscillation may impede the propagation of sequential information across nodes in a network. Together, these point to a potential role for a small number of neurons to play a larger role in relaying information, via oscillations, across several interconnected microcircuits.

Our general finding that spike-timing dependent plasticity (STDP) is a key mechanism to re-stabilize network dynamics provides a potentially new role for STDP in the injured brain. As a primary mechanism associated with Hebbian learning, STDP is typically considered as a mechanism to restructure the synaptic connections in a network after a training stimulus (51,52). The return of activity to a damaged network using STDP is reminiscent of how homeostatic plasticity allows a healthy network to gravitate towards a target activity rate (53). Interestingly, homeostatic plasticity may play a very different and destabilizing role in networks after either focal or more diffuse deafferentation of neurons (27), leading these networks into brief bursts of activity the resemble interictal discharges that appear in posttraumatic epilepsy (54,55). However, this rebalancing of networks with STDP has its limits, as the relative success of recovering initial dynamics is not complete at the highest injury levels. In light of several reports showing that one form of plasticity – long term potentiation (LTP) – is lost after traumatic injury *in vivo* and *in vitro* (56,57), our results emphasize the importance of therapeutic strategies to help promote plasticity after injury and regain initial network dynamics. Similarly, given that oscillations can be important to establish coherence among brain regions, steps to maintain plasticity in an injured network would likely improve network communications in the traumatically injured brain.

Our results also provide some suggestions on how local damage in the brain may affect both local and global brain dynamics. Cognitive disruptions from TBI are frequently viewed as changes in the network structure among regions in the brain (reviewed in (13,50,58–60). Commonly, past studies focus on the functional and structural deficits in the connections among nodes in the network resulting from diffuse axonal injury (29,50,61). With new techniques in medical imaging and network theory, we can begin to understand how changes in the connectivity between regions can impact higher level cognitive function (62–64). However, these approaches rely on maintaining function at the node (microcircuit) level after injury. Certainly, we know there are regions of the brain that can impart large cognitive deficits simply with their own malfunction (50,65). From the current work, we know that neurodegeneration can impact both network activity and neural oscillations in the node. In combination, our results suggest that damage to one node (microcircuit) could indirectly influence the coordination of activity across many connected brain areas.

It is certainly plausible to explore some of the unique consequences of local neurodegeneration in broader brain networks using oscillator or neural mass models to link structural and functional networks of the brain (66,67). Although these models capture gross behavior of network dynamics, the current formulation of these models lack the nodal accuracy to determine how perturbations caused by injury can impact the larger network function. Similar to work showing how local gamma activity could create biologically realistic BOLD correlations (32), changes in network oscillation frequency or oscillation width would likely have broader impact beyond the local network. Particularly important would be examining the implications of an inconsistent oscillatory rate, as seen in our model, in a connected oscillatory model. To our knowledge, this feature is not commonly explored in neural mass or oscillator-based models. However, there is evidence to suggest that these models have synchronization properties that are sensitive to nodal dynamic changes, but oscillators with individually variable frequencies have yet to be investigated (68,69). Similarly, both neural mass and oscillator-based models can potentially benefit from further analysis of biological plasticity mechanisms to repair from damage, given the role we found plasticity in stabilizing or practically recovering nodal dynamics.

Overall, this study indicates that neurodegeneration alters population-level activity and network oscillations, with subpopulation-dependent changes to oscillation frequency or duration. These changes in network dynamics can be significantly recovered with spike-timing dependent plasticity. We anticipate that future work brain network dynamics to specific patterns of damage will be valuable for gaining insight into specific patterns of damage causing longlasting alterations in brain dynamics, and other patterns of damage that may produce only temporary changes in neural dynamics. At a higher level, discriminating between these two injury patterns can help identify injuries that could cause lasting cognitive deficits much earlier than currently possible, pointing to an opportunity to treat and improve outcome in a vulnerable population of TBI survivors.

## Methods

### Modeling a representative cortical circuit

To investigate the connection between neural degeneration and functional network activity, we constructed computational neural networks of integrate-and-fire neurons ((33); Summary in Figure 6). Networks of 1000 neurons were constructed with 80% regular-spiking, excitatory neurons and 20% fast-spiking inhibitory to mimic the ratios commonly used to model cortical circuits (28,32,33). These neurons follow a system of ordinary differential equations to track membrane potential, membrane recovery, and threshold-based spiking as follows:

**Figure 6.**
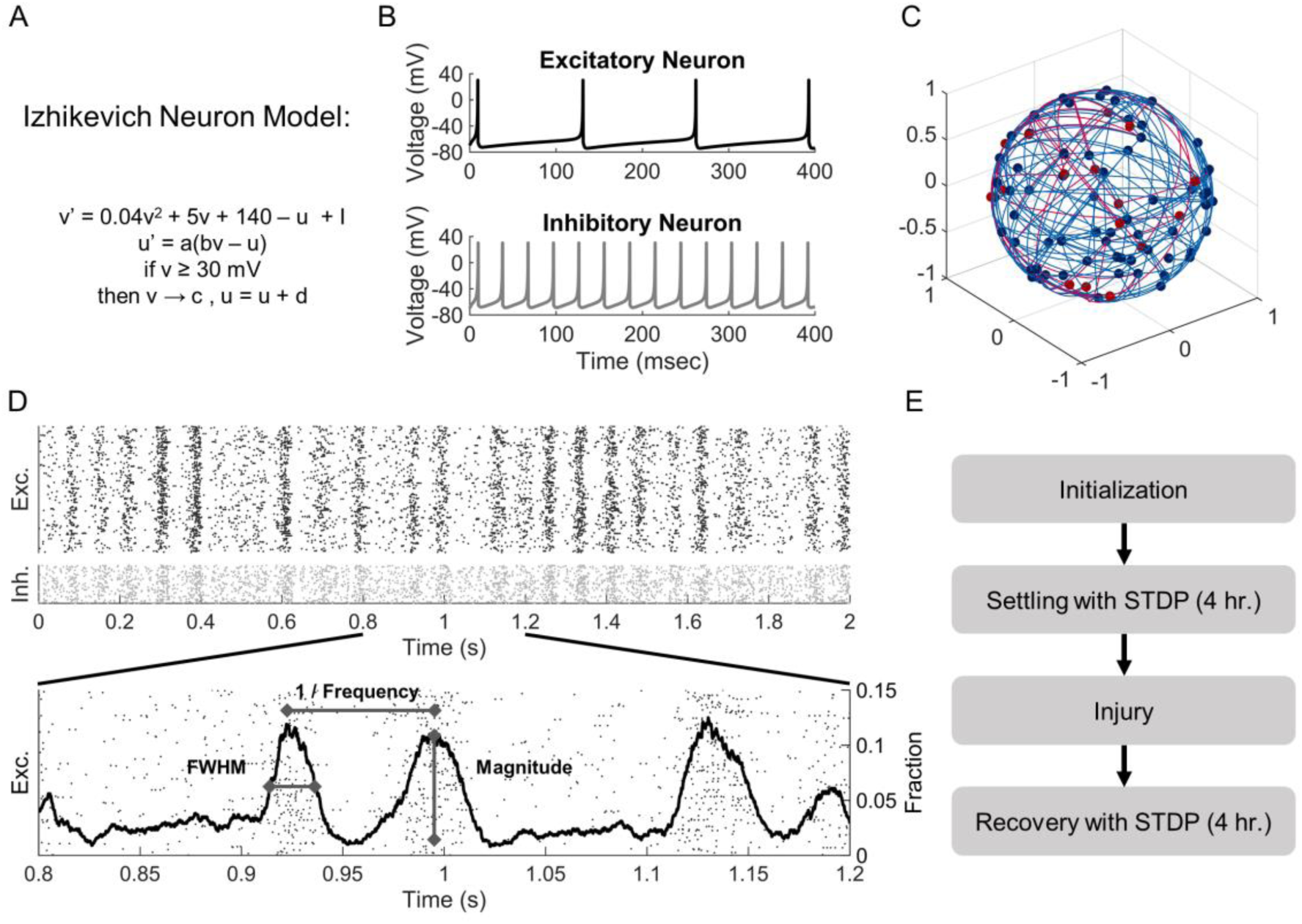
Schematic of Methods. **(A)** The Izhikevich integrate-and-fire neuron model was used to simulate neuron activity. **(B)** Model parameters were tuned to represent regular-spiking excitatory (Exc.) neurons and fast-spiking, low-threshold inhibitory (Inh.) interneurons. **(C)** Neurons were randomly placed on the surface of a sphere and connected into a 1000 node network. **(D)** Networks developed rhythmic activity oscillations after a four-hour period of connectivity weight settling with spike-timing-dependent-plasticity (STDP). **(E)** We characterized oscillations in our networks based on the number of oscillations per second, the peak number of spikes within our oscillation as a fraction of our network, and the full-width-half-maxium (FWHM). **(F)** After network weights settled with STDP, we injured networks at damage levels from 5% to 95%. After recording activity metrics immediately after injury, the networks restructured connectivity weights, again according to STDP. We then reassessed network function.

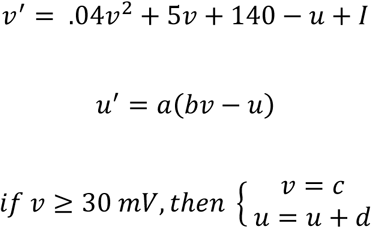

Where *v* represents the membrane potential in millivolts, and *u* is the membrane recovery variable. Parameters *a, b, c*, and *d* were set to create heterogeneous regular-spiking excitatory neurons and fast-spiking, low-threshold inhibitory neurons as in Izhikevich 2003 (Figure 6B). We allowed parameters *a, b, c*, and *d* to vary within a tight range for each neuron to avoid network behavior resulting from a homogeneous neuron population. We used a fixed timestep (0.2 millisecond) and a forward Euler method to compute *v* and *u* over time.

Consistent with average firing rates *in vivo* and previous models using Izhikevich neurons (28,35,36), we used a gamma distribution (*k*, θ = 2, 1/2) to randomly (f= 1 Hz) inject currents into individual neurons within the network. This stimulation was strong enough to cause the neuron to fire and send AMPA-or GABA-based synaptic signals to downstream targets. Synaptic currents were modeled as exponential decays from AMPA or GABA receptors, with τ = 5 msec. AMPA and GABA receptors were initially set to create EPSP/IPSP in line with past *in vivo* recordings (70). Repeated input stimuli to were attenuated at 40% immediately after a spike occurred (τ=150 msec) to model desensitization in the neuron population.

To avoid bias from edge effects in seeding neuron position, we placed neurons randomly on the surface of a unit sphere ((28); Figure 6C). The number of outputs for each neuron, which was drawn from a normal distribution with an average of 100 total outputs and inputs per neuron, varied slightly from each neuron to mimic features estimated in cortical circuits (10% variance; (71)). Neurons were randomly connected to each other across the surface, producing network properties of a classic Erdos-Renyi random graph (72). In a subset of simulations, we examined the effects of weak and strong distance-dependent connections and found that our main findings were unchanged. As a result, we present only the simulations using a random connection topology. Finally, we implemented synaptic transmission delays that were proportional to the distance between two neurons along the arclength of the sphere and set 8 ms as the maximum delay, consistent with *in vivo* recordings (73–77).

In neural networks, synaptic connection strengths adapt according to different models of synaptic plasticity. Among the models available, we chose to implement spike-timing dependent plasticity (STDP) because of its critical role in learning and the potential role this feature may play in cognitive deficits after traumatic injury (56,78,79). We used the Song model of STDP (34,80), in which the synaptic strength was adjusted based on the relative timing of synaptic inputs to a neuron and the subsequent action potential firing of the target neuron. Mathematically, this can be described as:

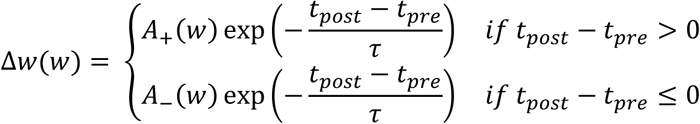

Where *W* is the synaptic weight of the connection between the pre- and postsynaptic neuron, *A*_*+*_ and *A*_*−*_, determines the maximum synaptic modification; *t*_*pre*_ and *t*_*post*_ are the timing of the pre- and postsynaptic activations; and *τ* is the plasticity time constant of 20 ms.

Given that this formulation of STDP leads to a bimodal distribution of synaptic weights, in which weights approach either the minimum or maximum possible weight (80), we seeded the synaptic strengths in our networks using a bimodal distribution. In addition, we restricted STDP to only effect synaptic weights between excitatory neurons in our model. For inhibitory neurons, we used a normal distribution of synaptic weights using a 10% variance. We scaled the synaptic weights of excitatory and inhibitory connections from the starting distributions [0 1] to [0 4] for excitatory neurons and to [-14 0] for inhibitory neurons. These scales were chosen to correspond to excitatory and inhibitory postsynaptic potentials recorded *in vivo* (70).

For each network, we constructed the network topology and assigned synaptic weights between connected neurons. We then allowed the network architecture to reweight with STDP until the firing behavior of neurons to achieve activity that did not vary in firing rate or average oscillation rate by more than 1% over a 5-minute simulation period. Across a range of connection architectures and synaptic weights, we achieved a stable activity pattern for simulation times of at least 4 hours. For each condition examined, we constructed and completed ten independent simulations, averaging the results from these simulations into a single group.

We recorded four measures from each simulation: the neuron-activity rate, oscillation frequency, oscillation peak magnitude, and oscillation width. Activity rate was calculated as the average firing rate of neurons within the network over a five-minute period. To evaluate the occurrence of oscillations, we recorded the number of action potentials occurring in a 10-millisecond sliding time window over the simulation time (Figure 6D). From these data, we selected times where peaks in activity occurred for the network (peak prominence ≥ 1) and used these times to compute the oscillation frequency as the number of oscillations per second. At each oscillation, we defined the oscillation magnitude as the maximum number of spikes within the sliding window and the width as the full width at half peak intensity.

We further analyzed each network oscillation to identify preferred spike timings for neurons. For each oscillation, we identified an interval that was 1.4 times the size of the oscillation width centered around the peak. We then split the interval into uniform deciles and assessed the likelihood for each neuron to fire within each time interval.

### Damaging the neural network

Random neuron injury: To mimic traumatic injury, neurons were randomly removed from the network after initial network reweighting with STDP. To maintain the initial excitatory-inhibitory balance of the circuit, 4 excitatory neurons were removed for every 1 inhibitory neuron. To assess the immediate effects of damage, we ran simulations without adjusting connectivity weights for 5 minutes, recording both the average firing rate and oscillation parameters over this time period. Next, we allowed networks to remodel with STDP with simulation times long enough to allow the firing rates and oscillation behavior to settle (4 hours). We then reassessed our metrics with a stable connectivity for an additional 5 minutes (Figure 6E).

Activity-based excitatory neuron injury: A second type of deletion scheme used the firing rate of individual neurons as the selection criteria for deletion. Once a given network restructured with STDP, we rank ordered the firing rate of each neuron within the network and deleted either the neurons with the lowest firing rate (LFR) or, alternatively, the neurons that displayed the highest firing rate (HFR).Because the focus of this removal strategy was to determine the effects of activity-based deletion, we opted to only remove excitatory neurons. Similar to random neuron injury, we assessed the immediate effects of neuron removal with a static connection weights for 5 minutes and then allowed the network to resettle for 4 hours.

### Statistical testing

To compare average activity and oscillation parameters between injury levels and baseline, we used one-way analysis of variance (ANOVA) with Tukey-Kramer post hoc test. Comparisons between injured and damaged networks at each injury level used t-tests with Bonferroni correction for multiple comparison.

